# Perineuronal nets on CA2 pyramidal cells and parvalbumin-expressing cells differentially regulate hippocampal dependent memory

**DOI:** 10.1101/2024.11.07.622463

**Authors:** Georgia M. Alexander, Viktoriya D. Nikolova, Tristan M. Stöber, Artiom Gruzdev, Sheryl S. Moy, Serena M. Dudek

**Affiliations:** Neurobiology Laboratory, National Institute of Environmental Health Sciences, Division of Intramural Research, National Institute of Health, Research Triangle Park, North Carolina 27713, USA; Carolina Institute for Developmental Disabilities and Department of Psychiatry, University of North Carolina School of Medicine, Chapel Hill, North Carolina 27599, USA; Institute for Neuroinformatics, Ruhr, University Bochum, Bochum, Germany; Department of Neurology, University Hospital Frankfurt, Frankfurt, Germany; Genome Editing and Mouse Model Core Facility, National Institute of Environmental Health Sciences, Division of Intramural Research, National Institute of Health, Research Triangle Park, North Carolina 27713, USA

## Abstract

Perineuronal nets (PNNs) are a specialized extracellular matrix that surround certain populations of neurons, including (inhibitory) parvalbumin (PV) expressing-interneurons throughout the brain and (excitatory) CA2 pyramidal neurons in hippocampus. PNNs are thought to regulate synaptic plasticity by stabilizing synapses and as such, could regulate learning and memory. Most often, PNN functions are queried using enzymatic degradation with chondroitinase, but that approach does not differentiate PNNs on CA2 neurons from those on adjacent PV cells. To disentangle the specific roles of PNNs on CA2 pyramidal cells and PV neurons in behavior, we generated conditional knockout mouse strains with the primary protein component of PNNs, aggrecan (*Acan*), deleted from either CA2 pyramidal cells (Amigo2 *Acan* KO) or from PV cells (PV *Acan* KO). Male and female animals of each strain were tested for social, fear, and spatial memory, as well as for reversal learning. We found that Amigo2 *Acan* KO animals, but not PV *Acan* KO animals, had impaired social memory and reversal learning. PV *Acan* KOs, but not Amigo2 *Acan* KOs had impaired contextual fear memory. These findings demonstrate independent roles for PNNs on each cell type in regulating hippocampal-dependent memory. We further investigated a potential mechanism of impaired social memory in the Amigo2 *Acan* KO animals and found reduced input to CA2 from the supramammillary nucleus (SuM), which signals social novelty. Additionally, Amigo2 *Acan* KOs lacked a social novelty-related local field potential response, suggesting that CA2 PNNs may coordinate functional SuM connections and associated physiological responses to social novelty.

**SIGNIFICANCE STATEMENT:** Perineuronal nets (PNNs) surround both inhibitory parvalbumin (PV)-expressing neurons and excitatory CA2 pyramidal neurons, but previous studies using enzymatic degradation cannot differentiate the relative roles of PNNs in the two populations. By conditionally deleting aggrecan (*Acan*) from CA2 pyramidal neurons without affecting PNNs on PV cells, and vice versa, we discovered distinct roles of PNNs on each cell type in behavior. Social memory, which requires CA2 activity, was impaired in mice lacking CA2 PNNs, but not in those lacking PV PNNs. Cognitive flexibility, assessed by reversal learning was also impaired in mice lacking CA2 PNNs, whereas contextual fear memory was impaired in those lacking PV PNNs. Thus, PNNs on each cell type differentially contribute to different forms of hippocampal-dependent memory.

## INTRODUCTION

Hippocampal area CA2 is a distinct part of the hippocampus owing to its molecular expression profile, synaptic properties and role in social memory (Dudek et al., 2016). Another unique feature of CA2 is the expression of perineuronal nets (PNNs) surrounding the excitatory pyramidal neurons there (Bruckner et al., 2003; Carstens et al., 2016). PNNs are aggregates of specialized extracellular matrix comprised of chondroitin sulfate proteoglycans that ensheathe select populations of neurons, primarily the parvalbumin (PV)-expressing inhibitory interneurons throughout the brain, and notably (excitatory) CA2 pyramidal neurons in hippocampus. Among their proposed functions, PNNs are thought to promote synaptic stabilization and innervation (Massey et al., 2006; Rowlands et al., 2018; Briones et al., 2020), as well as to negatively regulate synaptic plasticity (Pizzorusso et al., 2002; Carstens et al., 2016). In CA2, PNNs surround synapses in the pyramidal cells layer and stratum radiatum (Carstens et al., 2016), where inputs from various intrahippocampal and extrahippocamal areas form synapses (Kohara et al., 2014; Botcher et al., 2014; Chen et al., 2020, for example).

Neurons in area CA2 are now well-appreciated to function in social memory (Hitti and Siegelbaum, 2014; Stevenson and Caldwell, 2014; Smith et al., 2016), and PNNs in CA2 likely contribute to this function. Postnatal PNN development parallels the onset of social novelty recognition (Diethorn and Gould, 2023), and aberrant expression levels of PNNs are found in mouse models with impaired social memory, which in some cases, can be rescued by returning PNN expression levels to that seen in controls (Carstens et al., 2021b; Cope et al., 2022; Rey et al., 2022). In addition, inputs from the hypothalamic supramammillary nucleus (SuM), which signal social novelty to CA2 and regulate hippocampal theta oscillations associated with exposure to novelty, target the PNN- enriched pyramidal cell layer there (Chen et al., 2020, Kirk and McNaughton, 1991; Jeewajee et al., 2008). Area CA2 also contributes to other forms of hippocampal dependent memory. For example, contextual fear conditioning could be enhanced in animals with CA2 neuronal activity manipulated (Alexander et al., 2019). Also, spatial learning, tested in the Morris water maze (MWM), was enhanced in a knockout strain that, unusually, shows Schaffer collateral LTP in CA2 (Lee et al., 2010), and chronic silencing of CA2 impaired reversal learning in the MWM, as animals perseverated on the previous location of the platform (Lehr et al., 2023). These findings, along with the critical role of CA2 in social memory, led us to ask whether PNNs, specifically those on CA2 neurons, contribute to any of these forms of hippocampal-dependent memory.

The most commonly used approach for studying PNN function is through enzymatic degradation of the PNNs using chondroitinase (ChABC) (Carstens et al., 2016; Cope et al., 2022; Dominguez et al., 2019; Pizzorusso et al., 2002). However, ChABC does not differentiate between PNNs on CA2 pyramidal cells and those on PV-expressing interneurons. Even Cre-dependent viral expression of ChABC specifically in CA2 pyramidal cells has the effect of disrupting PNNs around PV cells, likely due to release of the enzyme into the neuropil (Carstens et al., 2021a). As our goal is to identify the specific role of PNNs on CA2 neurons distinct from those on PV-expressing neurons in hippocampal-dependent memory functions, an approach using ChABC is of limited use.

To overcome this limitation, we generated a mouse strain for deleting aggrecan (*Acan*), the primary chondroitin sulfate proteoglycan component of PNNs (Matthews et al., 2002), in a Cre-dependent way (floxed *Acan*). By breeding these mice with mice expressing Cre recombinase under control of either the *Amigo2* or *Pvalb* promoters, we deleted *Acan,* and thus PNNs, from either CA2 pyramidal cells or PV-expressing interneurons, respectively. We tested these two strains for social memory, spatial memory and fear memory. We found that PNNs on each cell type contribute to distinct forms of memory, with CA2 PNNs, but not PV PNNs, being important for social and spatial memory, specifically reversal learning, whereas PV PNNs, but not CA2 PNNs, were important for contextual fear memory. We also found that staining for vesicular glutamate transporter 2 (vGluT2), a marker for SuM inputs (Halasy et al., 2004) was reduced in Amigo2 *Acan* KOs, and a physiological local field potential (LFP) response to novel social investigation, a shift in the peak theta frequency, was not seen in Amigo2 *Acan* KOs. These findings clarify the specific role of PNNs on each of CA2 pyramidal cells and PV cells. Further, our findings suggest a functional relationship between CA2 PNNs, SuM inputs and an associated physiological response to recognition of social novelty.

## METHODS

### Animals

The *Acan* conditional null (flox) locus was generated by inserting loxP sites into intron 3- 4 and intron 4-5, placing the loxP sites approximately 10.5 kb apart so that following Cre- mediated recombination, excision of exon 4, which codes for a critical domain of ACAN protein, would occur. A single donor plasmid was constructed to introduce both loxP sites as a single gene editing event. The final donor plasmid consisted of (5’ to 3’): 639 bp 5’ homology arm (chr7: 78, 735, 306 - 78, 735, 944 GRCm39/mm39), unique BstBI restriction site, 5’ loxP site, internal homology arm including exon 4 and flanking intronic sequence (chr7: 78, 735, 955 - 78, 736, 229 GRCm39/mm39), unique SacII restriction site, 3’ loxP site, and 653 bp 3’ homology arm (chr7: 78, 736, 235 - 78, 736, 887 GRCm39/mm39). Gene targeting was done in B6129F1 embryonic stem cells (G4; 129S6/SvEvTacx C57BL/6Ncr). ES cells were lipofected with a 6:1 molar ratio of donor plasmids and Cas9-Puro/sgRNA (GCCCAAGGTCCCCTGGATCCTGG) delivery plasmids (pSpCas9(BB)-2A-Puro (PX459) V2.0), a gift from Feng Zhang (Addgene plasmid #62988; Ran et al., 2013). Following transfection, the cells were exposed to 48-hours of puromycin selection (0.9 µg/mL) followed by standard clonal expansion/screening. Clones were screened with long range PCR that included both the 5’ and 3’ homology arms and the genetic payload: Acan Flox Scr Fwd: ATCAGCAAGTGAGTTAAGTGGCA, Rev: TAGCTCTTCCTGAGGCCAATATG). The Cas9 target site is located in intron 3-4, so only clones that are homozygous for the distal 3’ loxP site in intron 4-5 were selected for downstream use. The closest potential off-target Cas9 site was 32 Mbp away in intergenic sequence, so no *in silico* predicted off-target sites were screened in the final *Acan* flox mouse line. Targeted ES cells were microinjected into albino B6J blastocysts for chimeric founder generation. The *Acan* flox allele was rescreened in F1 heterozygous mice. The line was then crossed to C57BL/6J wildtype mice to establish and expand the colony. The final mouse line was genotyped at Transnetyx (Cordova, TN) using primer/probe assays for the wild type, flox, and Cre-recombined null allele. To assess genetic background, the Neogen miniMUGA platform was run on three heterozygous floxed *Acan* flox mice via Transnetyx. Of the 10,819 nuclear SNPs on the miniMUGA platform, all mice were homozygous at 10,430 SNPs (96.4%), with 99.8% of those homozygous SNPs consistent with the C57BL/6J background. Of the remaining 389 SNPs, 316 SNPs had poor quality calls in all or some samples, while the other 73 variable SNPs were even distributed throughout the genome with no specific chromosomal hotspot, confirming that are final parental floxed *Acan* line was on the C57BL/6J genetic background.

Floxed *Acan* animals were bred with either Amigo2-cre animals (Alexander et al., 2017; McCann et al., 2021) or Pvalb-2A-Cre animals (Jax stock #012358; Madisen et al., 2010). Both strains are non-inducible, and for the Amigo2 line, Cre is expressed during postnatal development as well as in adulthood. Experiments were carried out in adult male and female mice of the resulting strains Amigo2-cre; *Acan* fl/fl (B6.Cg-Acan<tm1Ehs> Tg(Amigo2-cre)8Ehs; “Amigo2 *Acan* KO”) and PV-cre; *Acan* fl/fl (B6.Cg-Acan<tm1Ehs> Pvalb<tm1.1(cre)Aibs>; “PV *Acan* KO”). Mice were housed under a 12:12 light/dark cycle with access to food and water *ad libitum*. Mice used for behavior were group housed and were naïve to any treatment, procedure, or testing when beginning the experiments described here. Those used for electrophysiology were group-housed until the time of electrode implantation, after which they were singly housed. Males and females were housed in the same colony room. All behavioral assays were performed during the light cycle, between 8 AM and 3 PM. All procedures were approved by the NIEHS Animal Care and Use Committee and the Institutional Animal Care and Use Committee of the University of North Carolina and were in accordance with the National Institutes of Health guidelines for care and use of animals. Both males and females, as defined by the presence or absence of the Y chromosome, were used in all behavioral studies. A total of 201 mice of the Amigo2 *Acan* KO strain were used in these studies, including 48 animals (23 male, 25 female) for the three-chamber social assays, 39 animals (21 male, 18 female) for direct social interaction, 38 animals (18 male, 20 female) for MWM testing, 49 animals (23 male, 26 female) for fear conditioning, 10 animals (all male) for electrophysiology, and 17 animals (all male) for immunohistochemistry. A total of 211 mice of the PV *Acan* KO strain were used in these studies, including 53 animals (28 male, 25 female) for the three-chamber social assays, 36 animals (16 male, 20 female) for direct social interaction, 56 animals (29 male, 27 female) for MWM testing, 56 animals (28 male, 28 female) for fear conditioning and 10 animals (all male) for immunohistochemistry. All testing was conducted by experimenters blinded to mouse genotype.

### Immunohistochemistry

Animals used for immunohistochemistry were euthanized with Fatal Plus (sodium pentobarbital, 50 mg/mL; >100 mg/kg) and underwent transcardial perfusion with 4% paraformaldehyde. Brains were post-fixed in 4% paraformaldehyde overnight before being transferred to phosphate buffered saline (PBS). Brains were sectioned on a vibratome at 40 μm and stored in PBS with 0.1% sodium azide. For Acan, PCP4 and PV immunofluorescent staining, brain sections were rinsed in PBS then PBS with 0.1% triton (PBST), then blocked for 1 h in 5% normal goat serum/PBST. Sections were incubated overnight in blocking solution plus the biotinylated lectin *Wisteria floribunda* agglutinin (WFA, Vector labs, B-1355, 1:1000) and/or primary antibodies raised against Acan (Millipore, AB1031, 1:500, raised in rabbit), PCP4 (Invitrogen, PA5-52209, 1:500, raised in rabbit), and/or PV (Swant, PV27, 1:1000, raised in rabbit). After several rinses in PBST, sections were incubated in the secondary antibodies conjugated to fluorophores, including AlexaFluor goat anti-rabbit 568 (Invitrogen, A11011, 1:500) and streptavidin 488 (Invitrogen, S43454, 1:500). Finally, sections were washed in PBS and mounted with hardset mounting medium with DAPI (Vector Labs). For co-staining of vGluT2, PCP4 and WFA, sections were rinsed in 0.2% PBST, blocked overnight in 0.2% PBST with 3% NGS, washed and incubated for 48 hours in PBS with 3% NGS and primary antibodies, including vGluT2 (Millipore, AB225-1, 1:5000), PCP4 and WFA. Sections were then rinsed and incubated in AlexaFluor goat anti-guinea pig 568 (Invitrogen A11075, 1:500), AlexaFluor goat anti-rabbit 488 (Invitrogen, A11034, 1:500) and streptavidin 633 (Invitrogen, S21375, 1:500). Images of hippocampi were acquired on a Zeiss LSM 880 Airyscan confocal microscope. Fiji software (Schindelin et al., 2012) was used to quantify vGluT2 immunofluorescence using rectangular ROIs of 200 μm x 50 μm in the pyramidal cell layer of CA2, defined by PCP4 expression, or rectangular ROIs of 150 μm x 30 μm along the outer granule cell layer of dentate gyrus, identified by DAPI staining.

### Social approach in a 3-chamber test

The 3-chamber social test was used to assess sociability and preference for social novelty (Moy et al., 2004). The 3-chamber social testing apparatus was a rectangular, 3- chambered box fabricated from clear plexiglass (Fig. 2A*i*). Dividing walls had doorways allowing access into each chamber. An automated image tracking system (Ethovision, Noldus) was used to track animal position over the course of the assay. The procedure consisted of three 10-min phases: a habituation period, a test for sociability, and a test for social novelty preference. In the habituation period, the mouse was placed in the center chamber and allowed to explore the apparatus with the doorways into the two side chambers open. The mouse was then returned to the center chamber and doors to the side chambers closed. For the sociability phase, an unfamiliar, sex-matched adult C57BL/6J mouse (“Novel”) was placed within small plexiglass cage with holes (1 cm diameter) drilled in it to allow nose contact and positioned within one of the side chambers. An identical small plexiglass cage, with no mouse in it, was positioned in the chamber on the opposite side of the apparatus. The chamber doors were then opened, and the test animal was allowed to explore the three chambers. The amount of time the subject animal spent in proximity (5 cm) of each cage was measured as interaction time. The mouse was then returned to the center chamber with the doors closed to set up the social novelty phase. The previously-investigated animal remained in the same cage (“Familiar”), and a new novel animal was placed in the previously empty cage (“Novel”). The doors were then reopened, and the test animal was allowed access to all chambers. The amount of time that the test mouse spent in proximity (5 cm) of each cage was measured as the interaction time.

### Direct social interaction

The direct social interaction test was used to assess social memory using methods based on previous reports (Hitti and Siegelbaum, 2014; Kogan et al., 2000). Test animals were placed in a clean cage (45 cm L, 25 cm W, 21 cm H) within a tall-sided, open-topped, black plexiglass enclosure (56 cm L, 54 cm W, 77 cm H) under red light for 20 min to habituate to the context. After the habituation period, a C57BL/6J (4-5 weeks old) stimulus mouse of the same sex was then placed into the cage with the test animal, activity monitored for 5 min (trial 1), and then the stimulus mouse removed (Fig. 2B*i*). The total amount of time that the subject animal engaged in social interaction with the stimulus mouse (anogenital sniffing, nose-to-nose interaction and allogrooming of the stimulus mouse) was recorded. Social interactions driven by the stimulus mouse were not included in the recorded social interaction time. The stimulus animal was then returned to the home cage for a 1-hr intertrial interval. In trial 2, the stimulus mouse was reintroduced to the subject animal’s cage and social interaction time was again monitored and recorded as in trial 1.

### Morris water maze

The MWM was used to assess spatial and reversal learning, swimming ability, and vision (Morris, 1984; Moy et al., 2008). The water maze consisted of a large circular pool (diameter = 122 cm) partially filled with water (45 cm deep, 24-26° C, made opaque by the addition of non-toxic white poster paint), located in a room with numerous visual cues. The procedure involved three different phases: a visible platform test, acquisition in the hidden platform task, and a test for reversal learning. In the visible platform test each mouse was given 4 trials per day, across 3 days, to swim to an escape platform cued by a patterned cylinder extending above the surface of the water. For each trial, the mouse was placed in the pool at 1 of 4 possible locations (randomly ordered), and then given 60 sec to find the visible platform. If the mouse found the platform, the trial ended, and the animal was allowed to remain 10 sec on the platform before the next trial began. If the platform was not found, the mouse was placed on the platform for 10 sec, and then given the next trial. Measures were taken of latency to find the platform and swimming speed via an automated tracking system (Noldus Ethovision). Following the visible platform task, mice were tested for their ability to find a submerged, hidden escape platform (12 cm diameter). Each mouse was given 4 trials per day, with 1 min per trial, to swim to the hidden platform. Criterion for learning was an average group latency of 15 sec or less to locate the platform. Mice were tested until the group reached criterion, with a maximum of 9 days of testing. When the group reached criterion, mice were given a 1-min probe trial in the pool with the platform removed. Selective quadrant search was evaluated by measuring percent time spent in each quadrant, and number of crosses over the location where the platform (the target) had been placed during training, versus the corresponding areas in the other three quadrants. Following the acquisition phase, mice were tested for reversal learning, using the same procedure as described above. In this phase, the hidden platform was relocated to the opposite quadrant in the pool. As before, measures were taken of latency to find the platform. Once criterion was reached, the platform was removed from the pool, and mice underwent a probe trial to evaluate reversal learning.

### Fear conditioning

Mice were evaluated for learning and memory in a conditioned fear test in conditioning and testing chambers (Near-Infrared image tracking system, MED Associates, Burlington, VT) using previously reported methods (Alexander et al., 2019). The procedure had the following phases: training on day 1, a test for context-dependent learning on day 2, and a test for cue-dependent learning on day 3. Two weeks following the first tests, mice were given second tests for retention of contextual and cue learning (Fig. 6A). For the training phase on day 1, each mouse was placed in the test chamber, contained in a sound-attenuating box, and allowed to explore for 2 min. The mice were then exposed to a 30- sec tone (80 dB) that co-terminated with a 2-sec scrambled foot shock (0.4 mA). Mice received two additional shock-tone pairings, with 80 sec between each pairing. On day 2, mice were placed back into the original conditioning chamber for a test of contextual learning. Levels of freezing (immobility) were determined across a 5-min session. On day 3, mice were evaluated for associative learning to the auditory cue in another 5-min session. The conditioning chambers were modified using a plexiglass insert to change the wall and floor surface, and a novel odor (dilute vanilla flavoring) was added to the sound-attenuating box. Mice were placed in the modified chamber and allowed to explore. After 2 min, the acoustic stimulus (80 dB tone) was presented for a 3-min period. Levels of freezing before and during the stimulus were obtained by the image tracking system, and freezing during the tone was used as a measure of cued fear. Learning retention tests were conducted two weeks following the first tests. No significant differences were detected between males and females within either of the genotypes, so males and females were combined for analyses.

### *In vivo* electrophysiology

Methods for *in vivo* electrophysiology were similar to those previously reported (Alexander et al., 2018). Five Cre- and 5 Cre+ male Amigo2 *Acan* KO animals, (8-10 weeks old) were implanted with electrode arrays. The animals were anesthetized with ketamine (100 mg/kg, IP) and xylazine (7 mg/kg, IP), then placed in a stereotaxic apparatus. An incision was made in the scalp, and the skull was cleaned and dried. One ground screw (positioned approximately 4 mm posterior and 2 mm lateral to Bregma over the right hemisphere) and four anchors were secured to the skull. The electrode wire bundle was then lowered into a hole drill over dorsal CA2 (−1.95 mm AP, −2.3 mm ML, −1.9 DV mm relative to Bregma). Electrode wires were connected to a printed circuit board (San Francisco Circuits, San Mateo, CA), which was connected to a miniature connector (Omnetics Connector Corporation, Minneapolis, MN). Electrodes consisted of stainless steel wire (44 μm) with polyimide coating (Sandvik Group, Stockholm, Sweden). A total of 10 wires were bundled and lowered to the target region. Neural activity was transmitted via a 32-channel wireless 10× gain headstage (Triangle BioSystems International, Durham, NC) and was acquired using the Cerebus acquisition system (Blackrock Microsystems, Salt Lake City, UT). Continuous LFP data were band-pass filtered at 0.3– 500 Hz and stored at 1,000 Hz. Recordings were referenced to a silver wire connected to the ground screw. Behavior was recorded on video and time-synchronized to physiology recordings using Neuromotive Software (Blackrock Microsystems).

LFP recordings were made while animals investigated in three conditions: a novel context, a novel social stimulus and a familiar social stimulus (Fig. 3A). Animals were brought into the recording room ∼30 min before starting recordings, and red-light illumination was used throughout. First, animals investigated a novel context for 5 min (T maze dimensions: 45 cm L, 60 cm W, 13 cm H, with 7.5 cm W interior). The T maze was positioned within a larger open-topped plexiglass enclosure (56 cm L, 54 cm W, 77 cmH). The first min of time that the animal was in the T maze (a period of high locomotor activity) were excluded from analysis, and for the remaining data, periods when the animal was investigating but not running were used for analysis. Next, animals were moved to a clean cage (45 cm L, 25 cm W, 21 cm H) positioned within the larger enclosure and allowed to habituate for 30 min. A novel male C57BL/6J (4-5 weeks old) was then added to the cage for novel social investigation. Periods when the test animal was actively investigating the stimulus animal (anogenital sniffing, nose-to-nose interaction and allogrooming) were used for analysis. After 5 min, the novel animal was removed from the cage, and the test animal remained in the cage for 1 hr, after which the same stimulus animal (now-familiar) was reintroduced to the cage. Again, periods when the test animal actively investigated the stimulus animals were used for analysis.

### Classification of Navigational Strategies

Methods used here were similar to those previously reported (Lehr et al., 2023). Navigational strategies were categorized into spatial (directed search, corrected path, corrected path, direct path or perseverance) and non-spatial (thigmotaxis, circling, random path, scanning, or chaining) with Rtrack, a machine learning based water maze analysis package (Overall et al., 2020). The center of the circular pool was determined by calculating the midpoint between the maximum and minimum x and y coordinates of superimposed trajectories for every unique water maze configuration. The radius of the circle was computed as half of the maximum distance between the maximum and minimum x or y coordinates. The code used for search strategy classification is available at: https://github.com/tristanstoeber/alexander2024behavioral

### Experimental design and statistical analyses

Male and female animals of each genotype and each strain were used for each behavioral assay, and assays were run in the same order for all groups. Power analyses were performed to determine the number of animals required for all assays with the note that sociability, novel social recognition, MWM and fear conditioning assays were performed at the University of North Carolina School of Medicine, where animals were transported to before 4 weeks of age, which was often before genotyping results were obtained, so we could not be certain of the number of Cre+ and Cre- animals of each litter were transported. Any animal that did not complete all phases of each assay was removed from analysis for that assay. In addition, animals that showed floating behavior rather than swimming or did not complete the acquisition or reversal phases of the MWM were removed from analyses for that assay. A total of 11 of the Amigo2 *Acan* KO strain were excluded (2 male Cre-, 3 male Cre+, 6 female Cre-) and no PV *Acan* KOs were excluded from MWM analysis.

LFP data were Fourier transformed to generate power spectral densities using Matlab (Mathworks) software, and the frequency, between 6 and 12 Hz, corresponding to peak power was recorded as the peak theta frequency. For total theta power measurements, LFP was filtered between 6 and 12 Hz, and the area under the curve was obtained. Graphpad Prism 10 (Graphpad) was used for all statistical analyses. Two-factor grouped comparisons were analyzed using repeated measured (RM) two-way ANOVAs and Bonferroni multiple comparisons tests, with reported *p* values corrected for multiple comparisons. Single factor comparisons with two groups were analyzed using two-tailed unpaired t-tests, with or without Welch’s correction for unequal variance. Equality of variance was tested by F-tests. Behavioral data using both males and females were analyzed by three-way ANOVA, including sex, genotype and the assay-specific variable as factors. No main effects or interactions with sex were identified, so behavioral data presented in Figures 2, 4 and 6 included combined males and females. However, as sex-dependent trends were seen for social assays and the Morris water maze, data split by sex for those assays are shows in Extended Data Figures 2-1 and 4-1, respectively. For all analyses, significance (*p*) was set to <0.05. Data were tested for normality using D’Agostino and Pearson tests. Specific statistical tests used and the results of the tests are listed in the figure legends. For all graphs, mean and standard errors (SEM) are plotted, and dots in graphs represent individual animals/observations.

## Results

### Cell type-specific PNN deletion

In the hippocampus, PNNs surround inhibitory PV-expressing interneurons and excitatory CA2 pyramidal cells. To study the various roles of PNNs on each of these cell types selectively, we generated a conditional knockout (“floxed”) strain for the *Acan* allele allowing for Cre-dependent excision of exon 4, coding for a critical domain, similar to a previously reported conditional *Acan* knockout strain (Rowlands et al., 2018). Breeding with either Amigo2 Cre or PV Cre animals yielded animals with Acan and PNNs absent from either CA2 pyramidal neurons or PV interneurons, respectively, in Cre+ animals (Fig. 1). Specifically, CA2 pyramidal cells and presumptive PV cells immunostained for Acan in Cre- Amigo2 *Acan* KOs. However, Acan stain was absent from CA2 pyramidal cells in Cre+ Amigo2 *Acan* KOs, while presumptive PV cells retained Acan stain, demonstrating the cell-type selectivity of *Acan* deletion from CA2 cells over PV cells in Amigo2 *Acan* KOs (Fig. 1A). In addition, whereas Cre- animals showed PNN expression, as identified by WFA staining, surrounding CA2 pyramidal cells and PV cells, Cre+ Amigo2 *Acan* KOs showed PNN expression on PV cells but not CA2 pyramidal cells (Fig. 1B), and, conversely, Cre+ PV *Acan* KOs showed PNN expression surrounding CA2 pyramidal cells but not PV cells (Fig. 1C). This cell-type selectivity of Acan deletion, and thus PNNs, therefore permits side-by-side examination of behavioral roles of PNNs surrounding each cell type in isolation.

**Figure 1.**
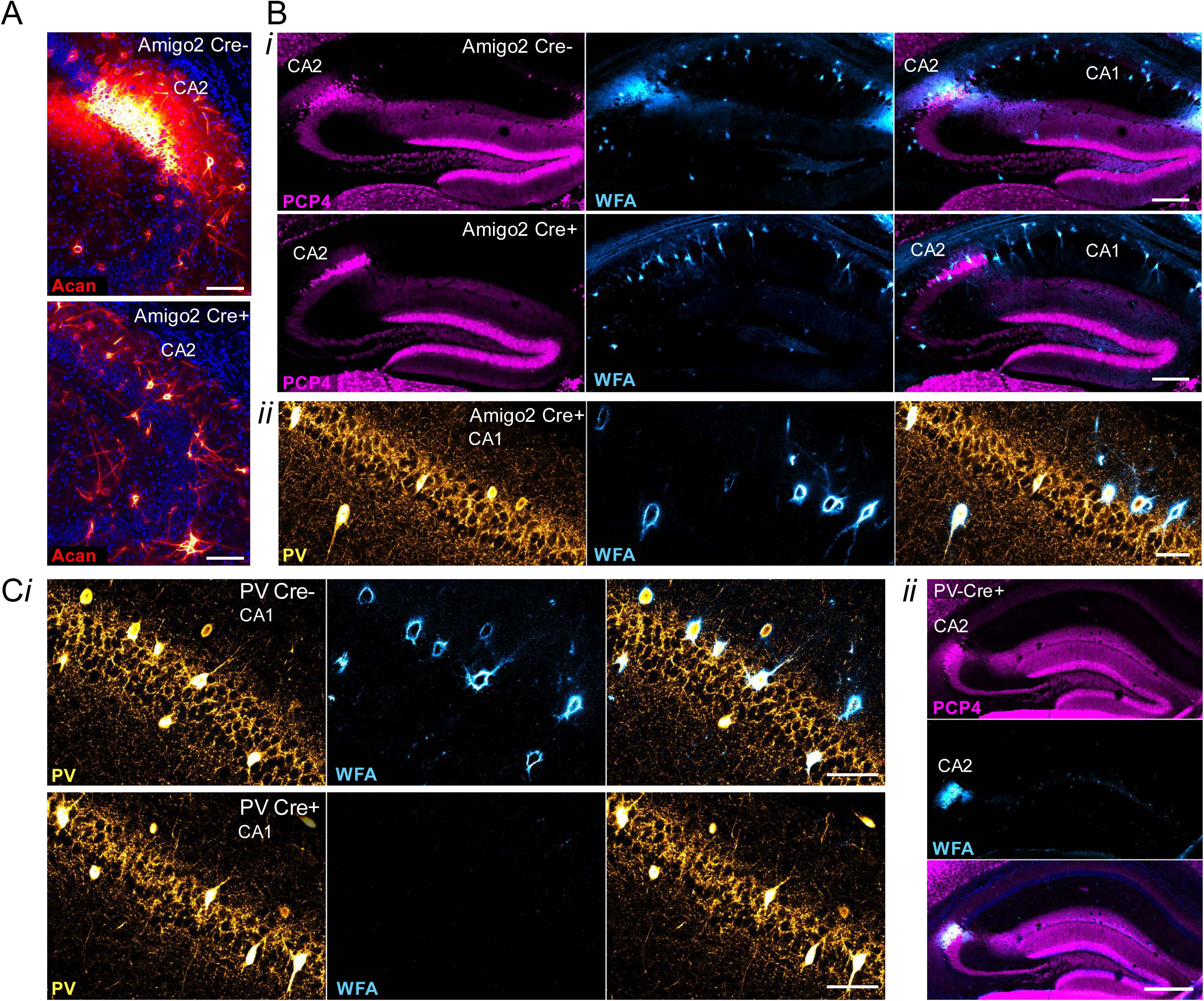
Cell-type specific *Acan* deletion results in a lack of Acan immunofluorescence and PNNs, as identified by WFA staining, around CA2 pyramidal cells or PV interneurons. A. Immunostaining for Acan shows the presence of Acan around CA2 pyramidal cells of Amigo2 Cre- tissue but the lack of such staining in Amigo2 Cre+ tissue. B. Cre+ Amigo2 *Acan* KO animals lack WFA stain on CA2 pyramidal cells (*i*) but retain WFA stain in PV-expressing cells (*ii*), whereas Cre- animals show normal WFA stain (*i*). C. Cre+ PV *Acan* KO animals lack WFA stain on PV-expressing cells (*i*) but retain expression on CA2 pyramidal cells (*ii*), as shown by colocalization with the CA2 marker, PCP4. Scale bars in A = 100 μm, in B = 250 μm (*i*) and 50 μm (*ii*), and in C = 50 μm (*i*) and 250 μm (*ii*).

### Amigo2 *Acan* KOs have impaired social memory

To determine whether loss of PNNs impacted social memory, Cre- and Cre+ Amigo2 and PV *Acan* KO mice were tested in each of three social assays, including a sociability assay and two assays of social recognition memory: preference for social novelty and direct social interaction. To determine whether sociability itself was impacted by PNN loss, we first tested whether animals showed a normal preference for a social stimulus over an inanimate object. In this assay, the amount of time a test animal spent interacting with a novel mouse in an enclosure compared with an empty enclosure was measured (Fig 2A*i*). Among the Amigo2 *Acan* KO animals, both Cre- control animals and Cre+ animals spent more time interacting with the novel animal than the empty enclosure, and the difference in interaction time with the two stimuli (social minus empty) was not significantly different between Cre- and Cre+ animals (Fig 2A*ii*). Similarly, among PV *Acan* KO animals, both control Cre- and Cre+ animals spent significantly more time interacting with the novel animal than the empty enclosure, and difference scores in interaction times were not significantly different between genotypes (Fig 2A*iii*). These data gave us confidence that social memory could be measured in subsequent tests.

Tests for social memory take advantage of the animals’ natural preference for social novelty. Here, we measured the amount of time the test animal spent in the interaction zone surrounding a novel animal or a familiar animal (Fig. 2B, Ext. Fig. 2-1A). For Amigo2 *Acan* KO animals, Cre- controls spent significantly more time in the interaction zone surrounding a novel animal than familiar animal. However, Cre+ Amigo2 *Acan* KO animals showed no significant difference in interaction time surrounding a novel versus familiar animal. The difference in interaction time with the two stimuli (novel minus familiar) was also significantly greater in Cre+ animals than Cre- animals (Fig. 2B*ii*). Among PV *Acan* KO animals, both Cre- controls and Cre+ KOs spent significantly more time with the novel animal than the familiar animal and difference scores were similar between the two genotypes (Fig. 2B*iii*). Therefore, deletion of PNNs from CA2 pyramidal cells, but not from PV cells, impairs preference for social novelty. These findings are typically interpreted as the mice having impaired memory for the social stimulus (Tan et al., 2019, for example).

**Figure 2.**
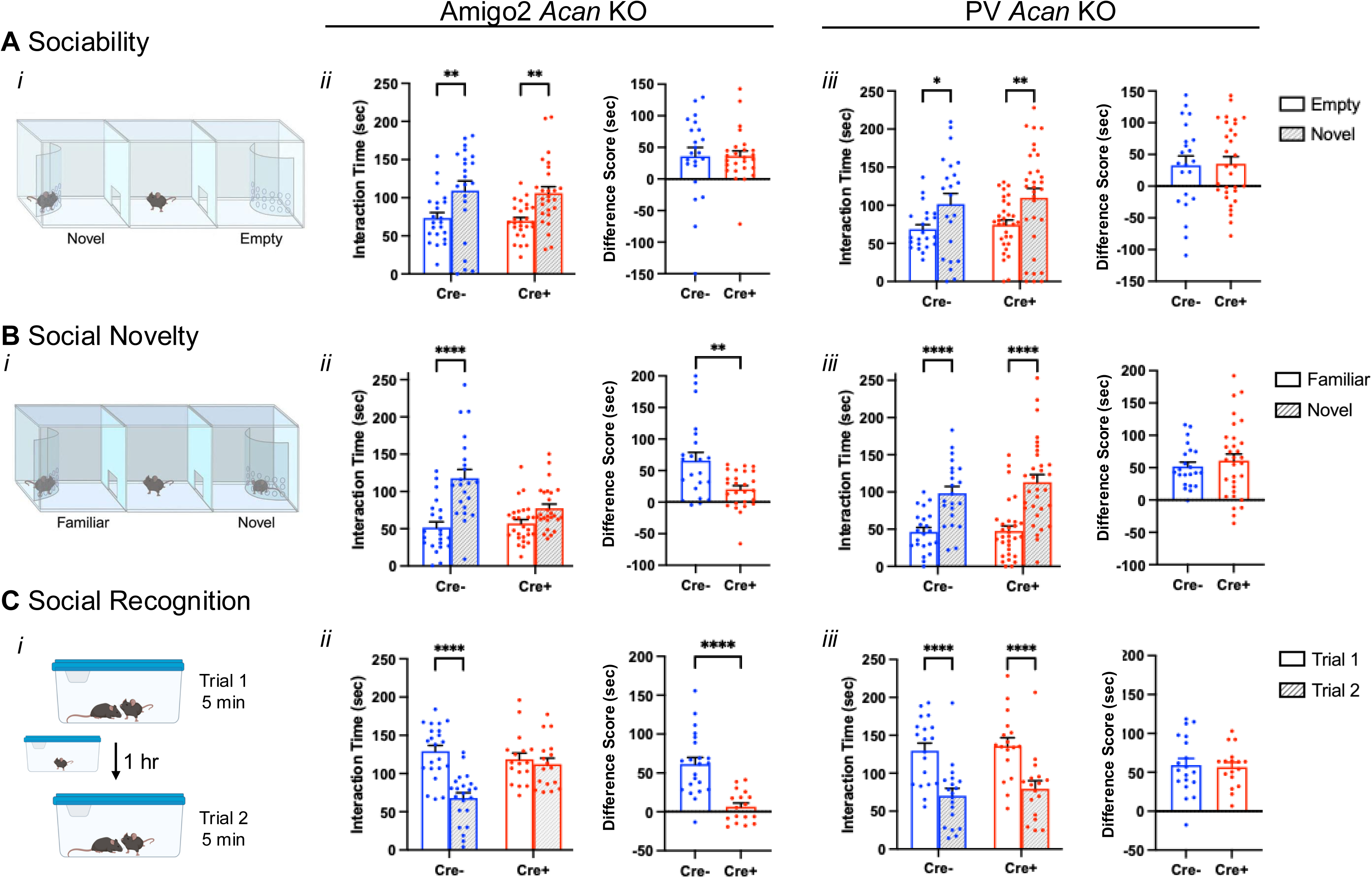
Amigo2 *Acan* KOs, but not PV *Acan* KOs, have impaired social recognition memory. A. In the sociability assay (*i*), animals can interact with a novel social stimulus within an enclosure or an inanimate empty enclosure, and the amount of time the animal spends within the interaction zone of each is measured. For each of the Amigo2 *Acan* KO (*ii*) and PV *Acan* KO (*iii*) strains, both Cre- and Cre+ animals showed a preference for interacting with the social stimulus (Amigo2 *Acan* KO: main effect of chamber: F(1,46)=21.86, *p*<0.0001, main effect of genotype: F(1,46)=0.16, *p*=0.70, interaction: F(1,46)=0.0013, *p*=0.97; multiple comparisons: Cre-: *p*=0.0057, Cre+: *p*=0.0022; PV *Acan* KO: chamber: F(1,51)=14.43, *p*=0.0004, genotype: F(1,51)=0.39, *p*=0.54, interaction: F(1,51)=0.018, *p*=0.89; multiple comparisons: Cre-: *p*=0.02, Cre+: *p*=0.0072), and difference scores (time interacting with novel social stimulus minus time interacting with the empty chamber) were similar between Cre- and Cre+ animals for each strain (Amigo2 *Acan* KO: t(33.9)=0.034, *p*=0.97; PV *Acan* KO: t(51)=0.14, *p*=0.89). B. In the preference for social novelty assay (*i*), animals have a choice between a novel and familiar mouse, and the amount of time the animal spends within the interaction zone of each is measured. Among Amigo2 *Acan* KO animals (*ii*), Cre- animals preferred the novel mouse but Cre+ animals did not (chamber: F(1,46)=41.10, *p*<0.0001, genotype: F(1,46)=4.29, *p*=0.044, interaction: F(1,46)=11.45, *p*=0.0015; multiple comparisons: Cre-: *p*<0.0001, Cre+: *p*=0.0605), and the difference score (novel minus familiar) was greater for Cre- than Cre+ animals (t(29.2)=3.21, *p*=0.0032). However, both Cre- and Cre+ PV *Acan* KO animals (*iii*) preferred the novel mouse (chamber: F(1,51)=72.23, *p<*0.0001, genotype: F(1,51)=0.64, *p*=0.43; multiple comparisons: Cre-: *p*<0.0001, Cre+: *p*<0.0001), and difference score was similar between genotypes (t(48.8)=0.73, *p*=0.47). C. In the direct interaction test of social recognition memory assay (*i*), animals were exposed to a novel animal in trial 1 and the same, now-familiar, animal in trial 2. For the Amigo2 *Acan* KO strain (*ii*) Cre- controls spent less time with the stimulus animal on trial 2 than trial 1, but Cre+ KOs spent equivalent time with the stimulus mouse in each trial (trial: F(1,37)=40.69, *p*<0.0001, genotype: F(1,37)=3.22, *p*=0.081; interaction: F(1,37)=20.69, *p*<0.0001; multiple comparisons: Cre-: *p*<0.0001, Cre+: *p*=0.85), and difference score (trial 1 minus trial 2) was significantly greater for Cre- than Cre+ animals (t(32.5)=5.58, *p*<0.0001). For the PV *Acan* KO strain (*iii*), both Cre- controls and Cre+ KOs, spent significantly less time with the stimulus animal on trial 2 than trial 1 (trial: F(1,34)=115.4, *p*<0.0001, genotype: F(1,34)=0.33, *p*=0.57; interaction: F(1,34)=0.064, *p*=0.80; multiple comparisons: Cre-: *p*<0.0001, Cre+: *p*<0.0001), and difference scores were similar between genotype (t(34)=0.25; *p*=0.80). Repeated measures two-way ANOVAs with Bonferroni multiple comparisons tests were used for all grouped comparisons, and results of multiple comparisons tests are shown on graphs. Two-tailed unpaired t-tests, with or without Welch’s correction for unequal variance, were used for difference score comparisons, and results are shown on graphs. \**p*<0.05, \*\**p*<0.01, \*\*\*\**p*<0.0001.

To further examine social recognition memory (Fig. 2C, Ext. Fig. 2-1B), we used a two-trial direct social interaction assay with a 1-hr intertrial interval. Typically, mice show a decrease in interaction time from trial 1 to trial 2, which is interpreted as memory for the previously investigated animal (Kogan et al., 2000). Here, the test animal was exposed to a novel animal for 5 min, and the same novel (now-familiar) animal for another 5 min after a 1-hr inter-trial interval, and the amount of time the test animal spent actively interacting with the stimulus animal was measured (Fig. 2C*i*). For the Amigo2 *Acan* KO strain, Cre- control animals spent significantly less time investigating the stimulus mouse on trial 2 than on trial 1, suggesting that the test animal recognizes the stimulus animal as familiar on trial 2. However, Cre+ Amigo2 *Acan* KOs did not show the expected decrease in interaction time with the stimulus animal on trial 2, suggesting that the Cre+ animals do not recognize the stimulus animal as familiar. In addition, the difference score in interaction time (trial 1 minus trial 2) was significantly greater for Cre- than Cre+ animals (Fig. 2C*ii*). By contrast, for the PV *Acan* KO strain, both Cre- controls and Cre+ animals showed a significant decrease in interaction time with the stimulus animal on trial 2 relative to trial 1 and difference scores did not vary by genotype (Fig. 2C*iii*). We interpret these findings to indicate that loss of CA2 PNNs causes impairment in an animal’s memory for a social stimulus.

### Novel social investigation increases peak theta frequency in control animals, but not in Amigo2 *Acan* KOs

To gain further insight into the nature of the social memory impairments seen in Amigo2 *Acan* KO animals, we recorded local field potentials (LFPs) from Cre- and Cre+ Amigo2 *Acan* KO animals during investigation of novel animals, presumably during the encoding of social memories. As a comparison, we first recorded LFPs during novel context investigation, in the absence of social stimuli. Finally, we asked how LFPs may differ upon investigation of novel and familiar social stimuli by recording LFPs upon re-exposure to the previously investigated animal (Fig. 3A-B). Animals were first placed in a novel context (T-maze) and allowed to explore for 5 min while LFPs were being recorded. Peak theta frequency during investigation of the novel context (measured during min 2-5 when animals were investigating but not running) did not differ between Cre- and Cre+ Amigo2 *Acan* KO animals (Cre- = 7.82 ± 0.07 Hz; Cre+ = 7.89 ± 0.036 Hz). During investigation of a novel animal, though, peak theta frequency increased significantly in Cre- control animals. However, this shift in peak frequency did not occur in Cre+ Amigo2 *Acan* KO animals (Cre- = 9.19 ± 0.14 Hz; Cre+ = 8.17 ± 0.11 Hz); peak theta frequency was significantly higher in Cre- animals compared with Cre+ animals (Fig. 3B*iii*). Upon subsequent re-exposure of Cre- animals to the (now-familiar) social stimulus, peak theta frequency significantly decreased from that seen upon novel social investigation but remained significantly greater than the peak theta frequency in the novel context. Peak theta frequency did not differ significantly between Cre- and Cre+ Amigo2 *Acan* KO animals upon familiar social investigation (Cre- = 8.63 ± 0.09 Hz; Cre+ = 8.24 ± 0.28 Hz). Area under the curve of the power spectral density in the theta range (6-12 Hz) was also collected as a measure of total theta power. Although total theta power was significantly influenced by stimulus within each genotype, no differences in total power were detected between genotypes.

**Figure 3.**
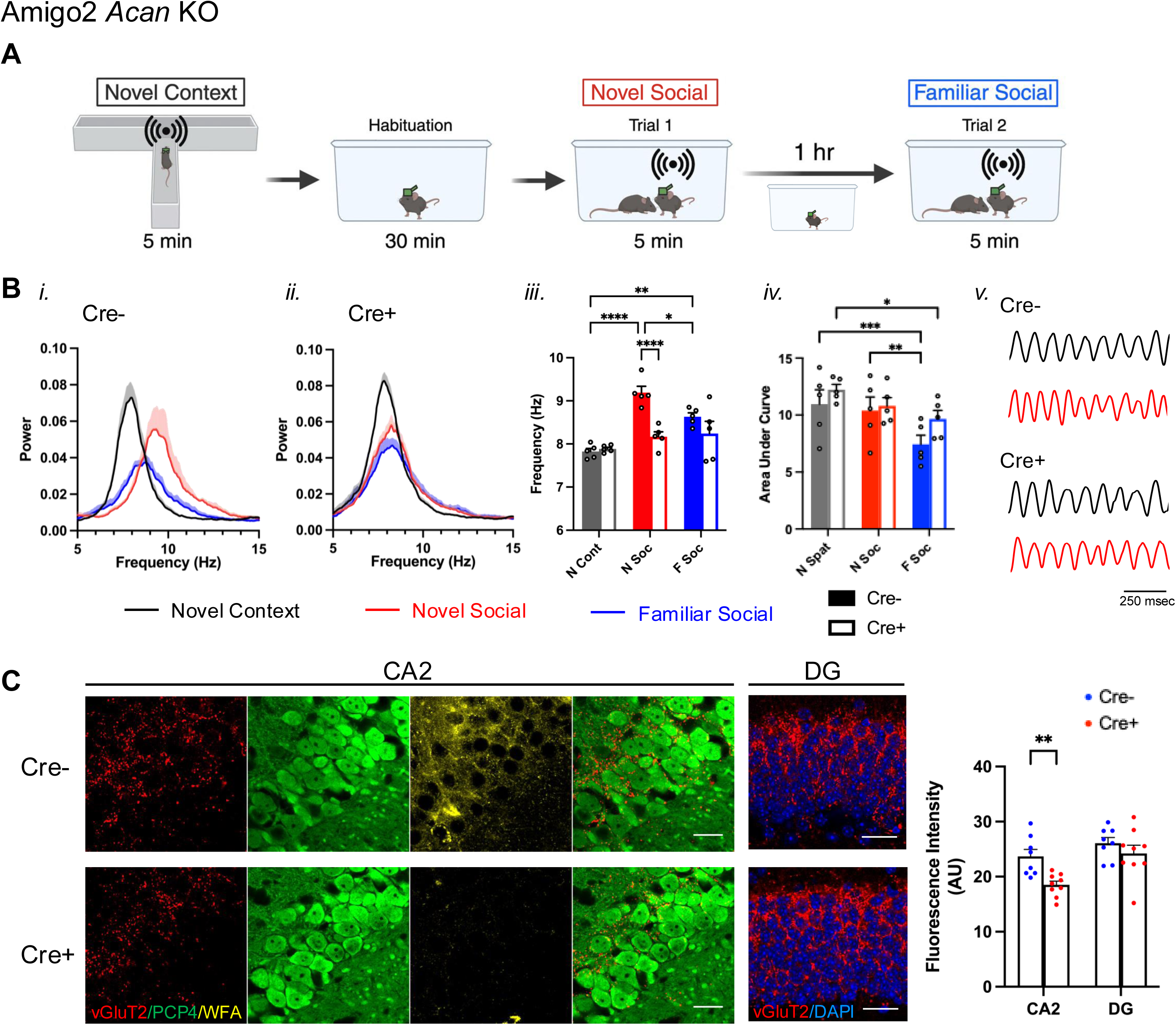
Peak theta frequency increases during novel social stimulus exploration in control animals but not Cre+ Amigo2 *Acan* KO animals, possibly due to impaired SuM inputs. A. LFP was recorded from CA2 while animals explored a novel context and were then habituated to a new clean cage before introducing a novel animal while recording LFP. One hour later, the same, now familiar, stimulus animal was reintroduced to the cage and LFP was recorded. B. Power spectral densities (*i-ii*), peak theta frequency (*iii*) and area under the curve of 6-12 Hz-filtered LFP power spectral density (*iv*) recorded from CA2 in Cre- (*i*), and Cre+ (*ii*) Amigo2 *Acan* KO animals during investigation in each condition. *iii*. In Cre- animals, peak theta frequency significantly increased during novel social investigation compared to the novel context, and then decreased upon re-exposure to the now-familiar stimulus animal but was still significantly increased compared to novel context exposure. By contrast, peak theta frequency was not significantly increased in Cre+ animals during novel social investigation (genotype: F(1,8)=12.77, *p*=0.0073; stimulus: F(2,16)=18.23, *p*<0.0001; interaction: F(2,16)=7.53, *p*=0.005). *iv*. Area under the curve of the theta-filtered power spectral density varied significantly across stimulus but did not differ between Cre- and Cre+ animals (genotype: F(1,8)=1.35, *p*=0.28; stimulus: F(2,16)=16.32, *p*=0.0001; interaction: F(2,16)=1.40, *p*=0.28). *v*. Representative theta-filtered (6-12 Hz) LFP traces recorded during investigation of a novel context (black) or a novel social stimulus (red). C. vGluT2, PCP4 and WFA co-staining of CA2 neurons shows vGluT2 positive terminals surrounding PCP4-expressing CA2 neurons, with WFA indicating the presence of PNNs in Cre- animals and the lack of WFA indicating the absence of PNNs around CA2 pyramidal cells of Cre+ animals. The red and green merged CA2 images show vGluT2 and PCP4 only. In the DG images, vGluT2 and DAPI only are shown. Measurements of vGluT2 fluorescent intensity from CA2 and DG reveal significantly less vGluT2 in CA2 of Cre+ Amigo2 *Acan* KO animals compared with Cre- control animals but no significant difference in DG (genotype: F(1,15)=6.29, *p*=0.024, subfield: F(1,15)=23.39, *p*=0.0002, interaction: F(1,15)=3.92, *p*=0.066). RM two-way ANOVA with Bonferroni multiple comparison tests used for all comparisons. Results of multiple comparisons tests shown on graphs. \**p*<0.05, \*\**p*<0.01, \*\*\**p*<0.001, \*\*\*\**p*<0.0001. Scale bars in C represent 25 μm.

PNNs are thought to function in synaptic stabilization, effectively holding synapses in place. In the absence of PNNs, synaptic targeting and stabilization may be impaired. One source of synaptic input to CA2 that is thought to play a central role in signaling social novelty is from the SuM. In addition, SuM is known to contribute to theta generation, and manipulation of SuM activity affects theta frequency (Kirk and McNaughton, 1991; Pan and McNaughton, 1997). Therefore, we labeled presumed synaptic terminals from SuM using immunofluorescence for vGluT2 (Halasy et al., 2004) and measured fluorescence intensity in the CA2 pyramidal cell layer. We found that vGluT2 immunofluorescence in Cre+ Amigo2 *Acan* KO animals was significantly less than in Cre- animals, suggesting decreased, or perhaps mistargeted, SuM inputs to CA2 (Fig. 3C). We also measured vGluT2 immunofluorescence in the dentate gyrus, where SuM also projects, and found no significant difference between Cre- and Cre+ Amigo2 *Acan* KO animals, suggesting that the impaired SuM inputs are selective to those projecting to CA2. We submit that SuM input to CA2 contributes to the increased peak theta frequency during novel social investigation, and decreased, or mistargeted, SuM input to CA2 may, at least partially, explain the lack of peak theta frequency increase in Cre+ animals during social investigation.

### Amigo2 *Acan* KO animals have impaired reversal learning in the Morris water maze

Prior work has shown that silencing the output of CA2 pyramidal neurons does not significantly impair spatial learning as assessed in a MWM, although the authors noted a trend toward the CA2 silenced animals learning more slowly, as measured by greater latency to reach the hidden platform on each day of acquisition training (Hitti and Siegelbaum, 2014). In contrast, knockout mice for RGS14, a protein enriched in CA2, showed enhanced learning in the MWM (Lee et al., 2010). Although reversal learning was not tested in RGS14 KOs, the CA2 silenced animals showed a trend toward an impairment in reversal learning (Hitti and Siegelbaum, 2014; Lehr et al., 2023). Therefore, we asked whether disruption of either CA2 PNNs (Fig. 4A, Fig. 4-1A) or PV PNNs (Fig. 4B, Fig. 4-1B) impacts spatial learning or reversal learning in the MWM test. For the Amigo2 *Acan* KO animals, during acquisition training, although overall Cre+ animals showed a greater latency than Cre- animals to reach the platform over training, both Cre- and Cre+ animals showed a significant decrease in latency to reach the platform from the first day to the last day of acquisition training. In addition, both Cre- and Cre+ animals spent significantly more time in the target quadrant than the opposite quadrant during the probe trial, and the difference score (percent of time in target quadrant minus percent of time in opposite quadrant) was similar between genotypes (Fig. 4A*i*).

**Figure 4.**
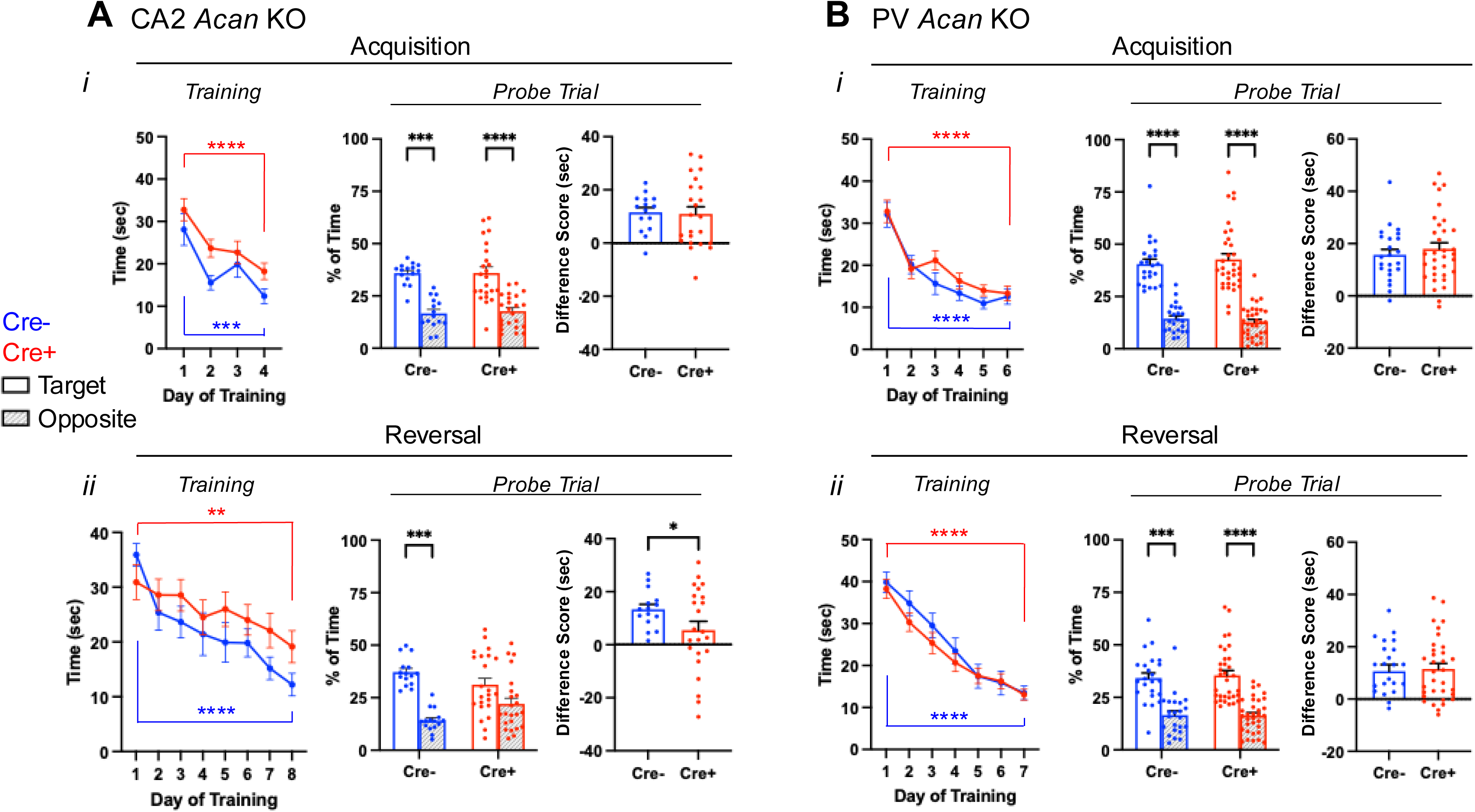
Morris water maze data for Amigo2 *Acan* KO (A) and PV *Acan* KO (B) animals. For each strain, data are split into that from the acquisition (top) and reversal (bottom) phases of the assay. For each phase, training data show the latency to reach the hidden platform over each day of training, and probe data show the percent of time that animals spent swimming in the quadrant of the pool where the platform was located during training and the opposite quadrant. Difference scores reflect the time in the target quadrant minus the time in the opposite quadrant. A*i*, In the acquisition phase, both Cre- and Cre+ Amigo2 *Acan* KO animals showed a significant reduction in latency to reach the platform from the first to the last day of training (training day: F(1,36)=39.87, *p*<0.0001; genotype: F(1,36)=3.54, *p*=0.07; interaction: F(1,36)=0.06, *p*=0.81). However, considering all days of training, Cre+ animals showed an overall greater latency to reach the platform than Cre- animals (training day: F(3,108)=14.62, *p*<0.0001; genotype: F(1,36)=5.93, *p*=0.02; interaction: F(3,108)=0.45, *p*=0.72). During the acquisition probe trial, both Cre- and Cre+ animals spent significantly more time in the target quadrant than the opposite quadrant (quadrant: F(1,36)=38.79, *p*<0.0001, genotype: F(1,36)=0.28, *p*=0.60; interaction: F(1,36)=0.027, *p*=0.87). The difference score (time in target quadrant minus time in opposite quadrant) was not significantly different between Cre- and Cre+ animals (t(35.3)=0.19, *p*=0.85). A*ii*. In the reversal phase of the assay, both Cre- and Cre+ animals showed a significant reduction in latency to reach the platform from the first day to the last day of training (training day: F(1,36)=35.84, *p*<0.0001; genotype: F(1,36)=0.11, *p*=0.74; interaction: F(1,36)=4.033, *p*=0.052), and no difference between genotypes was detected considering all days of reversal training (training day: F(7,252)=9.04, *p*<0.0001; genotype: F(1,36)=1.77, *p*=0.19; interaction: F(7,252)=1.15, *p*=0.33). However, on the reversal probe trial, whereas Cre- spent significantly more time in the target quadrant than the opposite quadrant, Cre+ animals did not (quadrant: F(1,36)=19.61, *p*<0.0001, genotype: F(1,36)=0.63, *p*=0.43; interaction: F(1,36)=3.64, *p*=0.065), and the difference score was significantly greater for Cre- animals than Cre+ animals (t(32.8)=2.06, *p*=0.047). B*i.* For PV *Acan* KO animals, in the acquisition phase, no differences were detected in latency to reach the platform from the first day to the last day of training (training day: F(1,54)=79.80, *p*<0.0001; genotype: F(1,54)=0.081, *p*=0.78; interaction: F(1,54)=0.000045, *p*=0.99), or over the entirety of training (training day: F(5,270)=31.30, *p*<0.0001; genotype: F(1,54)=1.12, *p*=0.29; interaction: F(5,270)=0.76, *p*=0.58). During the acquisition probe trial, both Cre- and Cre+ animals spent significantly more time in the target quadrant than the opposite quadrant (quadrant: F(1,54)=106.9, *p*<0.0001, genotype: F(1,54)=0.034, *p*=0.85; interaction: F(1,54)=0.48, *p*=0.49), and the difference score was not significantly different between Cre- and Cre+ animals (t(54)=0.69, *p*=0.49). B*ii.* In the reversal phase of the assay, both Cre- and Cre+ animals showed a significant reduction in latency to reach the platform from the first day to the last day of training (training day: F(1,54)=226.5, *p*<0.0001; genotype: F(1,54)=0.20, *p*=0.66; interaction: F(1,54)=0.12, *p*=0.74), and no difference between genotypes was detected considering all days of reversal training (training day: F(6,324)=61.70, *p*<0.0001; genotype: F(1,54)=0.55, *p*=0.46; interaction: F(6,324)=0.71, *p*=0.64). On the reversal probe trial, both Cre- and Cre+ animals spent significantly more time in the target quadrant than the opposite quadrant (quadrant: F(1,54)=44.52, *p*<0.0001; genotype: F(1,54)=0.29, *p*=0.59; interaction: F(1,54)=0.073, *p*=0.79) and the difference score was similar between genotypes (t(54)=0.27, *p*=0.79). RM two-way ANOVAs with Bonferroni multiple comparisons tests were used for all comparisons, and multiple comparison test results are shown on graphs. Two-tailed unpaired t-tests, with or without Welch’s correction for unequal variance, were used for difference score comparisons. \**p*<0.05, \*\**p*<0.01, \*\*\**p*<0.001, \*\*\*\**p*<0.0001.

Interestingly, in the reversal phase of the MWM, Cre+ Amigo2 *Acan* KO animals showed an impairment (Fig. 4A*ii*). Although both Cre- and Cre+ animals decreased latency to reach the platform from the first day to the last day of training, with no overall significant difference detected between genotype, on the probe trial Cre- animals spent significantly more time in the target quadrant than the opposite quadrant but Cre+ animals did not. In addition the difference score (target minus opposite quadrant) was significantly greater for Cre- animals than Cre+ animals, demonstrating an impairment in reversal learning among Amigo2 *Acan* KO animals. For PV *Acan* KO animals, no impairments were found in either the acquisition phase tests or reversal phase tests (Fig. 4B). Therefore, CA2 PNNs appear to contribute to reversal learning in the MWM test in a similar manner to that reported in CA2 silenced animals.

Of note, when data are split by sex, during the reversal training phase Cre+ males did not show a significant reduction in latency to reach the platform from the first day to the last day of training, although females did, and in the probe trial both male and female Cre+ animals failed to show a significant preference for the target quadrant (Fig. 4-1A). For PV Acan KO animals though, neither males nor females showed any impairment in the MWM (Fig. 4-1B).

### Amigo2 *Acan* KO animals use non-spatial search strategies in the Morris water maze

Given the similar MWM reversal learning findings in the Amigo2 *Acan* KO animals to those previously reported in CA2 silenced animals, we assessed another aspect of reversal learning that was also examined in the CA2 silenced animals: behavioral search strategies employed (Fig. 5; Lehr et al., 2023). Strategies used during each trial of each day of reversal leaning were categorized into various spatial and non-spatial forms using Rtrack, a machine learning based water maze analysis package (Overall et al., 2020). Males and females were analyzed separately because females showed improved performance over the course of reversal training but males did not (Fig. 4-1A*ii*), and aspects of spatial search strategies have been reported to differ according to sex (Zorzo et al., 2024). We found that from the first day to the last day of reversal training, both male and female control Cre- Amigo2 *Acan* KO animals significantly increased use of spatial strategies over time (Fig 5A*ii*, B*ii*). However, both male and female Cre+ animals did not significantly increase use of spatial strategies from the first to the last day of reversal training. These findings suggest that CA2 PNNs contribute to cognitive flexibility during spatial learning.

**Figure 5.**
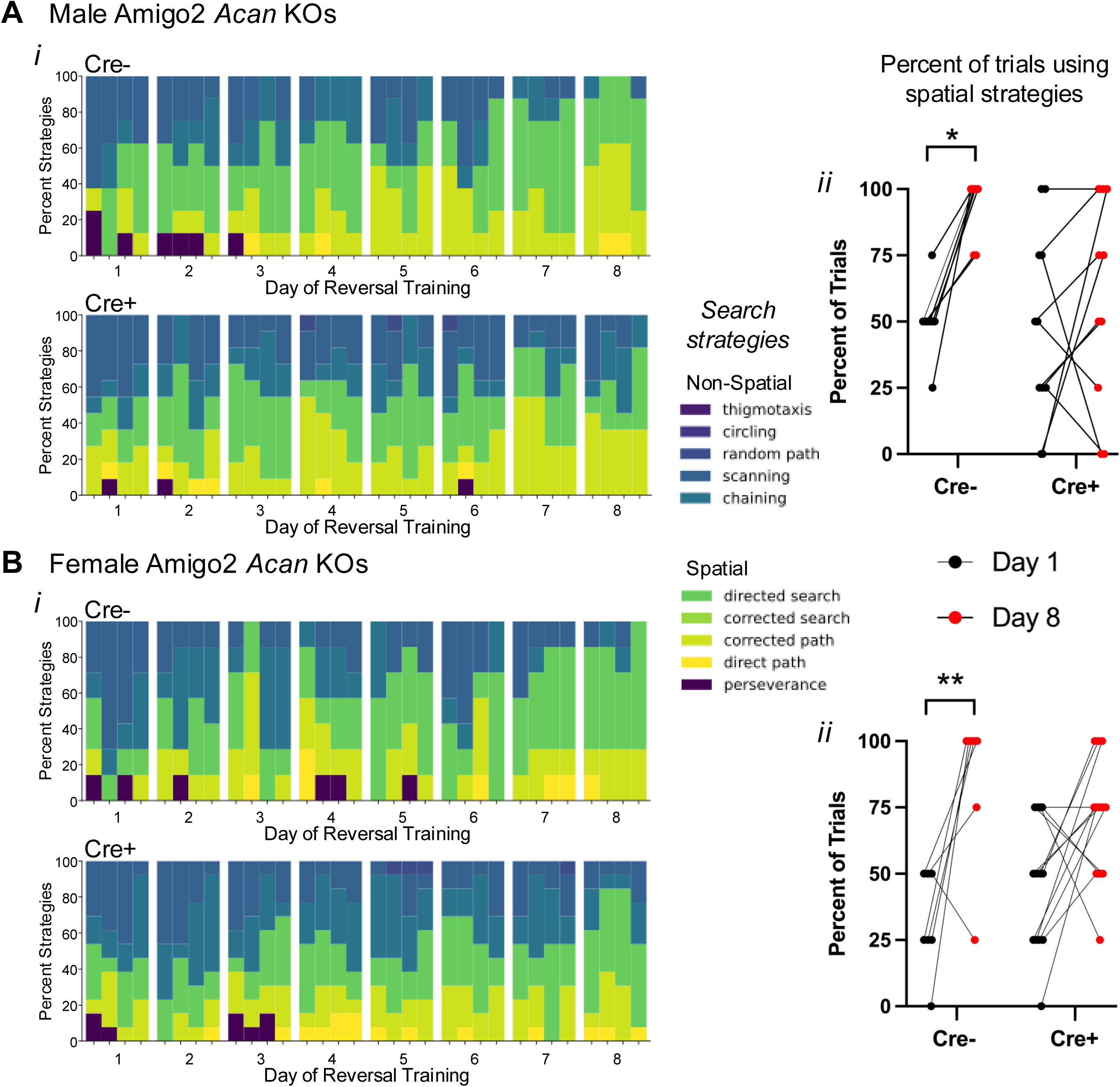
Cre+ Amigo2 *Acan* KOs persisted in using non-spatial search strategies during reversal learning training in the MWM among both males (A) and females (B). A*i*, B*i*. The search strategies used during each trial for each animal were categorized using Rtrack, a machine learning based water maze analysis package (Overall et al., 2020) (4 trials per day, 8 days of training) and plotted as the percent of animals using each search strategy for each trial of each day. Data are shown blocked into days and trials within each day. A*ii*, B*ii*. The percent of trials in which animals used spatial search strategies on the first day (day 1) and the last day (day 8) of reversal training were directly compared. Among males (A) and females (B), Cre- animals show a significant shift toward using spatial search strategies from day 1 to day 8, but Cre+ animals showed no significant change in the percent of trials using spatial search strategies (males: main effect of day: F(1,17)=10.31, *p*=0.0051, main effect of genotype F(1,17)=2.61, *p*=0.12; interaction: F(1,17)=2.84, *p*=0.11; multiple comparisons: Cre-: *p*=0.01, Cre+: *p*=0.51; females: main effect of day: F(1,18)=17.26, *p*=0.0006, main effect of genotype F(1,18)=0.0022, *p*=0.96, interaction: F(1,18)2.28, p=0.15; multiple comparisons: Cre-: *p*=0.005, Cre+: *p*=0.08). RM two-way ANOVAs with Bonferroni multiple comparisons tests were used for each sex. \**p*<0.05, \*\**p*<0.01.

### PV *Acan* KO animals have impaired contextual fear learning

Although CA2 is not known for its role in fear memory, one report described an effect of CA2 activity manipulation on contextual fear memory in females (Alexander et al., 2019). To determine whether PNNs in CA2 or PV neurons contribute to this type of non-social, non-spatial form of hippocampal-dependent memory, we tested Amigo2 *Acan* and PV *Acan* KO animals for contextual and cued associative learning using fear conditioning (Fig. 6). Animals were trained to associate a context and a tone cue with a shock stimulus, and on subsequent days freezing (immobility) was measured, in the absence of shock stimulus, when animals were returned to the context or placed in a new context and the tone played (Fig. 6A). For Amigo2 *Acan* KO animals, no differences were found in freezing for Cre+ animals compared to Cre- animals when returned to the context or played the cue tone (Fig. 6B). By contrast, Cre+ PV *Acan* KO animals showed an impairment in contextual fear learning. When freezing was measured in the context on day 2 of the assay (one day after shock pairing) as well as on day 16 (for memory retention testing), Cre+ PV *Acan* KO animals showed significantly less time freezing than Cre- animals. Time spent freezing did not differ by genotype in the PV *Acan* KOs in response to the cue. To account for multiple tests being run on each animal in the context and cue tests, data were entered into a three-way ANOVA with the factors of test type (context, cue), test number (test 1, test 2), and genotype (Cre-, Cre+), which revealed a significant interaction between genotype and test type that can be attributed to the genotype differences in contextual freezing in PV *Acan* KO animals. These findings suggest that PV PNNs play a role in context fear learning and/or memory. However, CA2 PNNs do not appear to be important for this type of learning.

**Figure 6.**
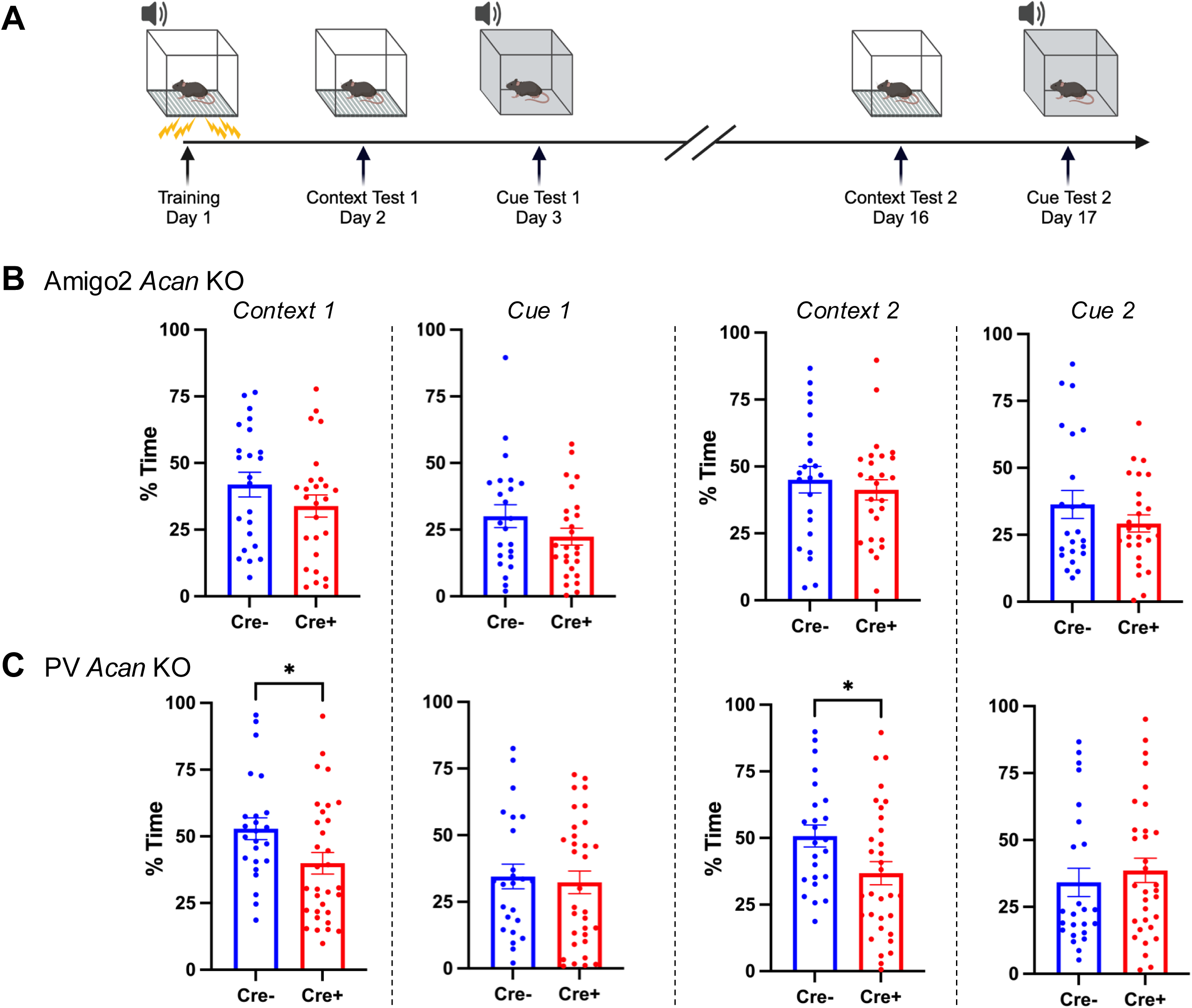
PV *Acan* KOs have impaired contextual fear learning. A. Animals were trained to associate a context and a tone with a shock stimulus on day 1. On subsequent days, animals were exposed to either the context or the tone in the absence of shock, and amount of time that the animal spent freezing was measured. B. Among Amigo2 *Acan* KOs, Cre+ animals did not differ from Cre- animals in percent of time freezing in either the context or in response to the cue when tested on days 2-3 or 16-17 of the assay (Context 1: t(47)=1.30, *p*=0.20; Cue 1: t(47)=1.47, *p*=0.15; Context 2: t(47)=0.62, *p*=0.54; Cue 2: t(47)=1.19, *p*=0.24). C. For the PV *Acan* KO animals, Cre+ animals showed significantly less freezing in response to the context compared with Cre- controls on each of day 2 and day 16 (Context 1: t(54)=2.22, *p*=0.03; Context 2: t(54)=2.27, *p*=0.027). However, no differences in freezing between Cre+ and Cre- PV *Acan* animals were detected in response to the cue (Cue 1: t(54)=0.34, *p*=0.73, Cue 2: t(54)=0.64, *p*=0.52). Two-tailed unpaired t-tests were used for all comparisons. \**p*<0.05.

## DISCUSSION

Hippocampal area CA2 is unique among the hippocampal subfields for several reasons, one of which is the presence of PNNs surrounding excitatory pyramidal neurons there. PNNs also surround inhibitory PV-expressing interneurons throughout brain, and within the hippocampus, intermingle with CA2 pyramidal neurons. Although much progress has been made using ChABC to study PNN function, it does not distinguish between PNNs on CA2 pyramidal neurons from those on PV expressing interneurons, limiting our ability to delineate the distinct functions of PNNs on each of these cell types. To overcome this limitation, we generated conditional knockout strains for *Acan*, the primary chondroitin sulfate proteoglycan of PNNs (Matthews et al., 2002), targeting either CA2 pyramidal neurons or PV-expressing interneurons, and assessed hippocampal dependent memory in each strain in parallel. We report that PNNs on CA2 pyramidal cells, but not on PV cells, contribute to social memory and reversal learning. Additionally, disruption of CA2 PNNs interferes with the normal input to CA2 from the SuM and with an electrophysiological signature of novel social exploration. Conversely, PNNs on PV cells, but not on CA2 pyramidal cells, contribute to contextual fear memory. Our findings support the conclusion that PNNs on CA2 neurons are integral to the functioning of CA2 neurons, which are increasingly appreciated to be active contributors to hippocampal function.

Previous findings have implicated PNNs as contributing to social memory. For example, the developmental onset of PNN expression in CA2 parallels the onset of preference for social novelty associated with social learning and memory (Diethorn and Gould, 2023), and direct injection of ChABC into hippocampus, targeting CA2, impairs social memory (Cope et al., 2022; Dominguez et al., 2019). Also, mouse models of Alzheimer’s disease, Rett syndrome, and autism spectrum disorder, all of which show impaired social memory, show aberrantly decreased or increased, respectively, staining for PNNs, and return of expression levels to that seen in control mice, can rescue the social memory impairments (Rey et al., 2022; Carstens et al., 2021b; Cope et al., 2022). These previously used approaches, although focusing on CA2, affect both PV cells and CA2 pyramidal cells. Using a genetic approach, we refined PNN deletion to either CA2 pyramidal cells or PV interneurons and show that PNNs on CA2 neurons, but not PV cells, are required for social novelty preference and social memory. We should note though, that taking the genetic approach, while permitting genetic access to control PNN deletion, does not provide the temporal control that ChABC does. Given that Cre is active in each strain before animals reach adulthood, we cannot rule out the possibility of compensatory changes between the time PNN expression begins (end of first postnatal week) to the time of testing (>8 weeks old). Additionally, as PV cells are found throughout the brain, PNN deletion on PV cells is not limited to those within hippocampus. These caveats aside, the genetic approach used here affords cell-type accuracy of PNN deletion not yet employed in the study of CA2, permitting study of PNNs on each cell type independently.

At least part of the mechanism by which CA2 engages in social memory likely involves SuM input. The developmental appearance of staining for the SuM axon terminal marker, vGluT2, parallels both the onset of preference for social novelty and the development of PNNs (Halasy et al., 2004; Carstens 2016; Diethorn and Gould, 2023). PNNs may promote synaptic innervation of CA2 neurons from SuM (Briones et al., 2020). They can direct axonal migration and promote stabilization through a variety of mechanisms such as by providing chemoattractant molecules like semaphorin 3A (Dick et al., 2013). PNNs are thought to act primarily as a matrix structure with holes that encircle synapses and are now recognized as a component of the tetrapartite synapse that includes the presynapse, postsynapse, astrocytic process and extracellular matrix structures (Ferrer-Ferrer and Dityatev, 2018; Sigal et al., 2019), placing PNNs in a critical location for synapse regulation. Indeed, PNNs surround synapses in the CA2 pyramidal cells layer and on dendrites (Carstens et al., 2016), areas that are contacted by intrahippocampal sources, as well as by the SuM (Botcher et al., 2014; Kohara et al., 2014; Chen et al., 2020). We found decreased vGluT2 staining in CA2, but not dentate gyrus, in animals lacking PNNs on CA2 neurons, suggesting reduced SuM input and/or mistargeting in CA2. The SuM is thought to signal social novelty through its direct projection to CA2 because, for example, exposure to a novel social stimulus increases cFos expression and neuronal activity in the SuM, and optogenetic activation of SuM terminals in CA2 increases interaction with a familiar social stimulus to levels seen upon exposure to a novel social stimulus (Chen et al., 2020). Also likely is that inputs from local interneurons that provide strong feed forward inhibition are mistargeted, as they also form synapses within the stratum pyramidale, collaborating with SuM inputs to likely shape spike timing that would contribute to memory formation (Robert et al., 2021).

To build on the social memory findings, we performed *in vivo* electrophysiological recordings while animals investigated a novel context, a novel animal, or a familiar animal. We focused on hippocampal theta oscillations given the role of SuM in regulating theta frequency and contribution of theta frequency to memory encoding (Kirk and McNaughton, 1991; Quirk et al., 2021) We found that whereas control animals showed an increase in peak theta frequency during novel social investigation relative to novel context investigation, Cre+ Amigo2 *Acan* KO animals did not show such a theta frequency shift, implicating impaired novelty signaling between SuM and CA2 neurons. The SuM is intimately involved in theta oscillations through its direct projection to medial septum (MS), which is required for hippocampal theta oscillations, as well as its direct inputs to the hippocampus (Vertes, 1992; Kocsis and Vertes, 1994; Haglund et al., 1984). In addition, the SuM appears to control theta frequency because SuM neurons fire at theta frequency and silencing SuM decreases hippocampal theta frequency (Kirk and McNaughton, 1991; Pan and McNaughton, 1997). Further, silencing SuM and thus decreasing theta frequency significantly impairs spatial learning (Pan and McNaughton, 1997), demonstrating a critical relationship between SuM firing, theta frequency, and learning. Social novelty increases firing rate of a subset of SuM neurons, specifically those that project to CA2 (Chen et al., 2020), but it is unknown whether those same SuM neurons also project to MS. CA2 pyramidal cells themselves project to the MS (Cui et al., 2013), so it is plausible that during novel social investigation, SuM inputs to CA2, likely in concert with inputs from local interneurons also targeted by SuM (Robert et al., 2021), drive and shape CA2 input to regulate MS activity and theta frequency. That we did not see an increase in theta frequency during novel social investigation in animals lacking PNNs in CA2 suggests that the absence of PNNs there impairs functional communication between SuM and CA2, to possibly impair communication with downstream CA2 targets, including MS. The decrease in vGluT2 staining in CA2, but not in DG, in the Amigo2 *Acan* KO mice, supports the idea that PNNs play a role in the development or maintenance of SuM inputs.

In the MWM test of spatial memory, both strains of *Acan* KOs were able to acquire the initial location of the hidden platform, suggesting intact ability to form a spatial representation of the environment and navigate to the platform using that representation. However, reversal learning was impaired in Cre+ Amigo2 *Acan* KO mice. The reversal phase of the MWM requires the animal to first recognize that the task has changed and then to form a new representation of the platform location relative to spatial cues. CA2 pyramidal cells are responsive to changes in environmental stimuli; they are place cells that encode space through firing patterns, and they remap in response to social or novel stimuli added to the environment (Alexander et al., 2016; Mankin et al., 2015; Oliva et al., 2016). This apparent responsivity of CA2 place cells to change suggests that they may actively contribute to the neural computations required for learning a new platform location during the reversal learning phase. Interestingly, when separated by sex (Ext. Fig. 4-1A), we only saw impaired reversal learning over the course of training in the males, suggesting that females lacking CA2 PNNs may be less dependent on PNNs or employ compensatory mechanisms to reduce the impact of lacking PNNs during training. That said, both males and females were impaired in the probe trial following reversal training, so reversal memory impairment is present in both sexes.

Analysis of behavioral search strategies used by Amigo2 *Acan* KOs during reversal training revealed that Cre- animals begin reversal training using non-spatial search strategies, and over the course of training, transition to spatially-guided strategies, suggesting successful encoding of a new spatial representation and possibly repression of the old one. By contrast, the animals lacking CA2 PNNs persist in using non-spatial search strategies over the course of reversal learning period, which one may speculate reflects impaired responsivity and encoding of CA2 neurons to the change in platform location. Interestingly, animals with CA2 chronically silenced similarly show an increased reliance on non-spatial search strategies over the course of reversal training compared with controls (Lehr et al., 2023).

Similar behavioral impairments in the MWM were also observed in mice lacking the mineralocorticoid receptor (MR; *Nr3c2*) in forebrain excitatory neurons (Berger et al., 2006). Within the brain, MR is most heavily expressed in the hippocampus, and within hippocampus, MR is highest in CA2 (Herman et al., 1989; Kalman and Spencer, 2002; McCann et al., 2021). Mice lacking MR also lack many of the molecularly defining features of CA2, including PNNs, and vGluT2 staining (McCann et al., 2021). This finding again supports the likely importance of PNNs, SuM, and CA2 as a whole in the adaptations required for reversal learning. However, a balance of PNNs appears to be necessary to proper CA2 functioning because similar behavioral impairments to those we’ve seen in Amigo2 *Acan* KOs are seen in BTBR mice, which have an overabundance of PNNs in CA2 and impaired social memory (Moy et al., 2007; Cope et al., 2022). Although BTBR mice acquire the location of the hidden platform during the acquisition phase, they too, show impaired performance in reversal learning in the MWM (Moy et al., 2007). Thus, PNN balance likely contributes to CA2 neuronal activity level and/or precision, which may impact their ability to adjust and encode a new target location using spatial cues.

Finally, PV *Acan* deletion, but not Amigo2 *Acan* deletion, impaired contextual fear learning. We previously demonstrated that manipulating CA2 neuronal activity increased freezing in shock associated context, an effect present in only females (Alexander et al., 2019). Although we may not expect to find an impairment in fear conditioning among males lacking CA2 PNNs here based on our previous finding, we were somewhat surprised that the females lacking CA2 PNNs were not impaired. That said, as described for reversal learning, the females may be less susceptible to impairment when lacking CA2 PNNs, so perhaps the same hold true for conditioned fear. By contrast, we did find an impairment in contextual fear memory in animals lacking PNNs on PV cells. Similarly, ChABC injections affecting all of hippocampus, or targeting CA1, have also been shown to impair contextual fear memory (Hylin et al., 2013; Liu et al., 2023). Thus, these previously reported conditioned fear memory impairments are likely attributable to loss of PNNs on PV cells and not CA2 pyramidal cells.

In summary, PNNs on CA2 neurons and PV cells differentially contribute to different types of hippocampal dependent memory, with those on CA2 pyramidal cells contributing to social memory and reversal learning and those on PV cells contributing to contextual fear memory. These findings build on previous reports of hippocampal PNNs in hippocampal-dependent memory to clarify the specific roles of PNNs on each of CA2 pyramidal cells and PV cells that could not be appreciated by enzymatic degradation of PNNs.

## Acknowledgements

We thank the expert staff at both the NIEHS Fluorescence Microscopy and Imaging Center and the NIEHS animal care staff for all their support.

## Extended Data Figure Legends

**Extended Data Figure 2-1.**
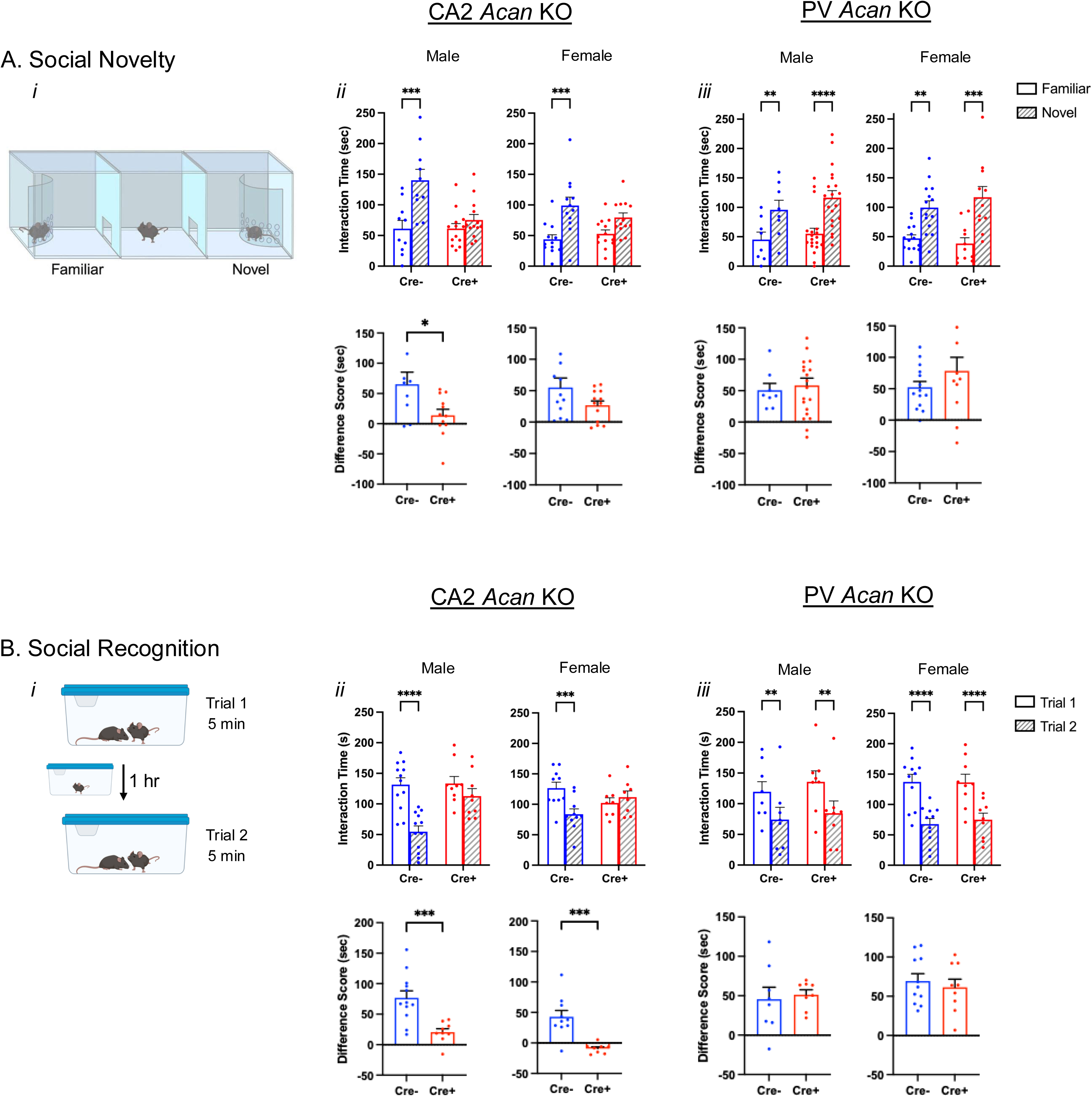
Amigo2 *Acan* KOs, but not PV *Acan* KOs, have impaired social recognition memory. A. In the preference for social novelty assay (*i*), animals have a choice between a novel and familiar mouse. Among CA2 *Acan* KO animals (*ii*), for both males and females, Cre- animals preferred the novel mouse but Cre+ animals did not (males: main effect of chamber: F(1,21)=17.6, *p*=0.0004, main effect of genotype: F(1,21)=6.3, *p*=0.02, interaction: F(1,21)=8.5, *p*=0.008; post hoc: Cre-: *p*=0.0002, Cre+: *p*=0.60; females: main effect of chamber: F(1,23)=26.01, *p*<0.0001, main effect of genotype: F(1,23)=0.28, *p*=0.60, interaction: F(1,23)=3.16, *p*=0.088; multiple comparisons: Cre-: *p*=0.0002, Cre+: *p*=0.050). Difference scores (novel minus familiar) showed that Cre- males preferred the novel mouse significantly more than Cre+ males, whereas difference scores were not significantly different for females (males: t(19)=2.49, *p*=0.022; females: t(23)=1.78, *p*=0.089). However, both Cre- controls and Cre+ PV *Acan* KO animals preferred the novel mouse. Preference for the novel was seen in both males and females (males: main effect of chamber: F(1,25)=33.69, *p<*0.0001, main effect of genotype: F(1,25)=0.89, *p*=0.35; females: main effect of chamber: F(1,23)=36.25, *p*<0.0001, main effect of genotype: F(1,23)=0.12, *p*=0.72). Difference scores were similar between the two genotypes for both males and females (males: t(26)=0.40, *p*=0.69; females: t(23)=1.21, *p*=0.24). B. In the direct interaction test of social recognition memory assay (*i*), animals were exposed to a novel animal in trial 1 and the same, now-familiar, animal in trial 2. For the CA2 *Acan* KO strain (*ii*), for both males and females, Cre- controls spent less time with the stimulus animal on trial 2 than trial 1, but Cre+ KOs spent equivalent time with the stimulus mouse in each trial (males: main effect of trial: F(1,19)=46.75, *p*<0.0001, main effect of genotype: F(1,19)=4.51, *p*=0.047; interaction: F(1,19)=15.55, *p*=0.0009; post hoc tests: Cre-: *p*<0.0001, Cre+: *p*=0.14; females: main effect of trial: F(1,16)=7.8, *p*=0.013; main effect of genotype: F(1,16)=0.022, *p*=0.88, interaction: F(1,16)=19.17, *p*=0.0005; post hoc tests: Cre-: *p*=0.0001, Cre+: *p*=0.61). Difference scores (trial 1 minus trial 2) were significantly greater for Cre- animals than Cre+ animals for each sex (males: t(19)=3.94, p=0.0009; females: t(16)=4.37, p=0.0005). For the PV *Acan* KO strain (*iii*), both Cre- controls and Cre+ KOs, of both the male and female sex, spent significantly less time with the stimulus animal on trial 2 than trial 1 (males: main effect of trial: F(1,14)=34.73, *p*<0.0001, main effect of genotype: F(1,14)=0.28, *p*=0.61; females: main effect of trial: F(1,18)=87.78, *p*<0.0001, main effect of genotype: F(1,18)=0.048, *p*=0.82). Difference scores were similar between genotypes (males: t(14)=0.35, *p*=0.73; females: t(18)=0.58, *p*=0.57). RM two-way ANOVAs with Bonferroni post hoc tests were used for all two-factor comparisons, and two-tailed unpaired t-tests were used for difference scores. Multiple comparisons test results are shown on graphs, \**p*<0.05, \*\**p*<0.01, \*\*\**p*<0.001, \*\*\*\**p*<0.0001.

**Extended Data Figure 4-1.**
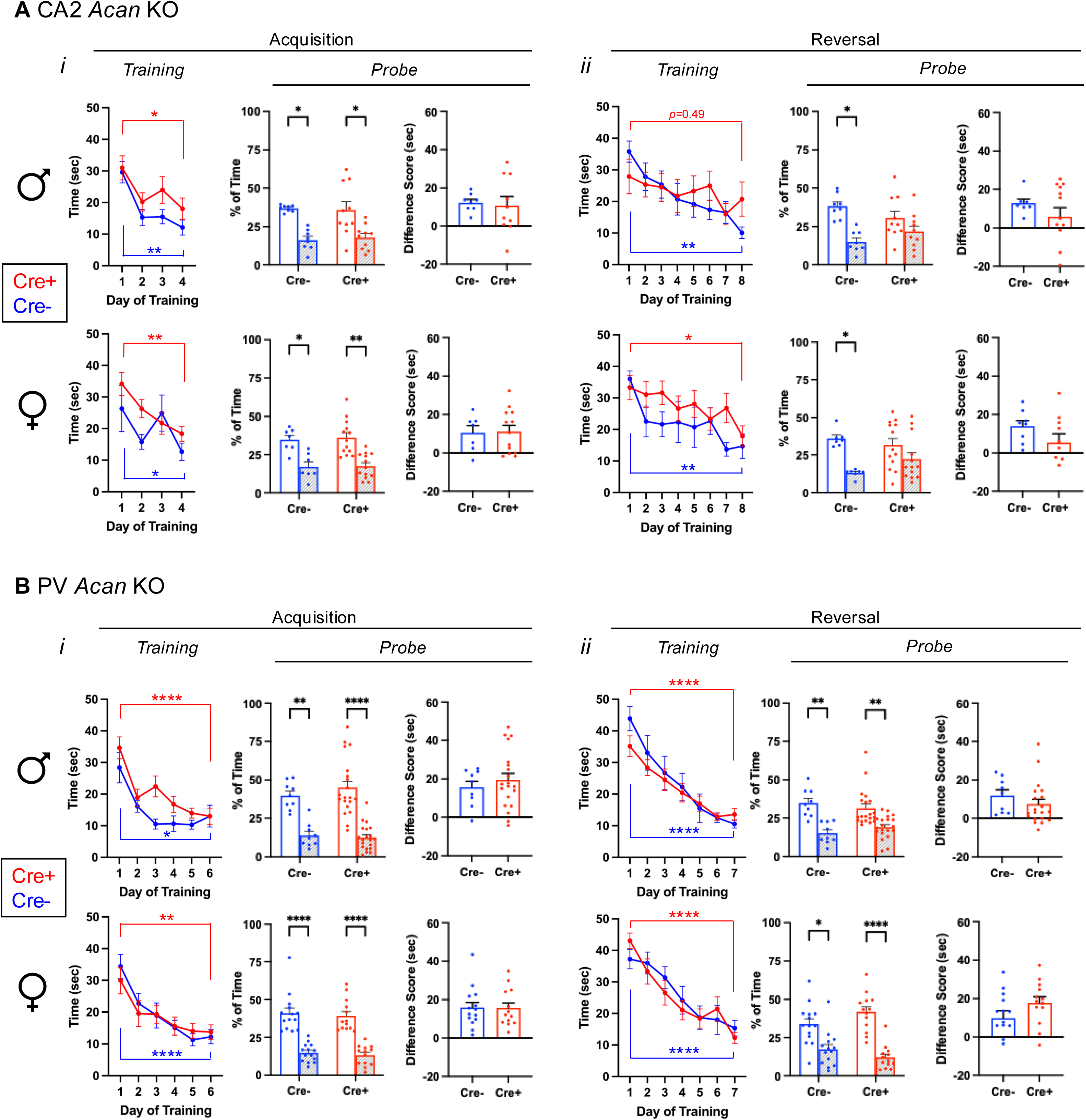
Morris water maze data for Amigo2 *Acan* KO (A) and PV *Acan* KO (B) animals. For each strain, data are split into that from males (top row) and females (bottom row) for the acquisition and reversal phases of the assay. For each phase, training data show the latency to reach the hidden platform over each day of training, and probe data show the percent of time that animals spent swimming in the quadrant of the pool where the platform was located during training and the opposite quadrant. Difference scores represent the time in the target quadrant minus time in the opposite quadrant. A*i*, For Amigo2 *Acan* KO animals, both males and females showed no difference between Cre- and Cre+ animals in latency to reach the platform during acquisition training (males: main effect of training day: F(3,48)=8.51, *p*=0.0001, main effect of genotype: F(1,16)=4.16, *p*=0.058; females: training day: F(3,54)=6.2, *p*=0.0011, genotype: F(1,18)=2.02, *p*=0.17), and all groups showed a significant reduction in latency to reach the platform from the first to the last day of training (males: Cre-: *p*=0.0044, Cre+: *p*=0.017; females: Cre-: *p*=0.033, Cre+: *p*=0.0039). Both Cre- and Cre+ animals also spent significantly more time in the target quadrant than the opposite quadrant during the acquisition probe trial (males: quadrant: F(1,16)=17.80, *p*=0.0007, genotype: F(1,16)=0.020, *p*=0.89; multiple comparisons: Cre- *p*=0.016, Cre+ *p*=0.019; females: quadrant: F(1,18)=18.24, *p*=0.0005, genotype: F(1,18)=0.6179, *p*=0.44; multiple comparisons: Cre- *p*=0.038, Cre+ *p*=0.0033). Difference scores were similar between genotypes within each sex (males: t(11.2)=0.33, *p*=0.75; females: t(18)=0.11, *p*=0.92). A*ii*. In the reversal phase of the assay, for both males and females, a main effect of training day was detected (males: training day: F(7,112)=4.91, *p*<0.0001; genotype: F(1,16)=0.11, *p*=0.74; females: training day: F(7,126)=3.96, *p*=0.0006; genotype: F(1,18)=2.08, *p*=0.17). However, multiple comparisons analysis revealed that whereas Cre- males had a significantly lower latency to reach the platform on day 8 compared to day 1 (*p=*0.0026), Cre+ males did not show a significant reduction in latency from day 1 to day 8 (*p=*0.49), suggesting an impairment in reversal learning among Cre+ male Amigo2 *Acan* KOs. Among females, both Cre- and Cre+ animals had a significantly reduced latency to reach the platform on day 8 compared to day 1 (Cre-: *p*=0.0087, Cre+: *p*=0.010). On the reversal probe trial, whereas Cre- males and Cre- females spent significantly more time in the target quadrant than the opposite quadrant, neither Cre+ males nor Cre+ females spent significantly more time in the target quadrant (males: quadrant: F(1,16)=10.39, *p*=0.0053, genotype: F(1,16)=0.14, *p*=0.72, interaction: F(1,16)=2.07, *p*=0.17; multiple comparisons: Cre-: *p*=0.013, Cre+: *p*=0.39; females: quadrant: F(1,18)=8.35, *p*=0.0098, genotype: F(1,18)=1.76, *p*=0.20, multiple comparisons: Cre-: *p*=0.041, Cre+: *p*=0.35). However, difference scores were not significantly different between genotypes for either sex (males: t(16)=1.44, *p*=0.17; females: t(16.21)=1.35, *p*=0.20). B*i.* For PV *Acan* KO animals, both males and females showed no difference between Cre- and Cre+ animals in latency to reach the platform during acquisition training (males: main effect of training day: F(5,135)=14.35, *p<*0.0001, main effect of genotype: F(1,27)=3.01, *p*=0.094; females: main effect of training day: F(5,125)=13.75, *p<*0.0001, main effect of genotype: F(1,25)=0.028, *p*=0.87). Both Cre- and Cre+ animals also spent significantly more time in the target quadrant than the opposite quadrant during the acquisition probe trial (males: quadrant: F(1,27)=40.97, *p<*0.0001, genotype: F(1,27)=0.57, *p*=0.46; females: quadrant: F(1,25)=63.45, *p<*0.0001, genotype: F(1,25)=1.10, *p*=0.30). Difference scores were similar between genotypes (males: t(27)=0.72, *p*=0.48; females: t(25)=0.041, *p*=0.97). B*ii.* In the reversal phase of the assay, for both males and females, Cre+ animals did not differ from Cre- animals in acquisition training (males: training day: F(6,162)=31.66, *p*<0.0001; genotype: F(1,27)=0.29, *p*=0.60; females: training day: F(6,150)=32.13, *p*<0.0001, genotype: F(1,25)=0.029, *p*=0.87). On the reversal probe trial, among males and females, both Cre- and Cre+ animals spent significantly more time in the target quadrant than the opposite quadrant (males: quadrant: F(1,27)=20.74, *p*=0.0001, genotype: F(1,27)=0.06, *p*=0.80, females: quadrant: F(1,25)=30.90, *p<*0.0001, genotype: F(1,25)=0.64, *p*=0.43). Difference scores were similar between genotypes (males: t(27)=1.06, *p*=0.3; females: t(25)=1.61, *p*=0.12). RM two-way ANOVAs with Bonferroni multiple comparisons tests were used for all two-factor comparisons, and two-tailed unpaired t-tests were used for difference scores. Multiple comparisons test results are shown on graphs. \**p*<0.05, \*\**p*<0.01, \*\*\*\**p*<0.0001.

